# Mitotic bypass and continued endocycling promote cancer cell survival after genotoxic chemotherapy

**DOI:** 10.1101/2025.11.03.686258

**Authors:** Kevin Truskowski, Louis Rolle, George Butler, Margaret E Yang, Kenneth J Pienta, Sarah R Amend

## Abstract

Genotoxic chemotherapies are central components of the treatment regimen for most cancers but are rarely curative. Drug-tolerant persister cells (DTPs) evade cell death during these treatments by accessing transient adaptive states and can contribute to cancer progression after treatment. Here, we demonstrate that cancer cells can survive genotoxic chemotherapy-induced stress by accessing a DTP state wherein stress-induced bypass of mitosis precipitates continued endocycling which promotes survival by allowing cells to evade mitotic catastrophe and cell death. Mechanistic studies indicate that persistent DNA damage signaling in endocycling persister cells triggers sustained p53-independent CDK1 inhibition by WEE1 and Myt1. Continued survival in endocycling persisters is dependent on activation of this G2 checkpoint, and disrupting WEE1 or Myt1 activity using clinical-stage small molecule inhibitors is sufficient to drive CDK1 reactivation, forcing mitotic entry, catastrophe, and cell death. Our results define endocycling DTPs as targetable mediators of cancer cell persistence after genotoxic therapy.

## Introduction

Genotoxic chemotherapies are central tools for the treatment of most cancers. These drug regimens commonly induce initial tumor regression but leave behind surviving populations that can seed relapse. Understanding which cancer cells survive treatment and how they do so remain urgent questions for the field. Resistance to chemotherapy has largely been attributed to models of genetic heterogeneity whereby a subpopulation of cells harboring a preexisting genetic advantage is selected for under therapeutic pressure, expands, and becomes the dominant population in the recurrent tumor. While evidence supporting this model is plentiful, it has become clear that non-genetic mechanisms of resistance are also common^1^. In lieu of genetic advantage, drug-tolerant persister cells (DTPs) survive drug treatment by accessing transient adaptive states and are increasingly recognized drivers of cancer lethality^2–4^. DTPs have been described in various cancer models, including lung, pancreatic, prostate, breast, and melanoma, and in response to both targeted therapies (e.g., kinase inhibitors) and cytotoxic drugs (e.g., platinum-based chemotherapy)^3^. Cancer cells’ rapid adaptation to diverse stressors highlights the significance of these states in determining treatment outcome.

Chemotherapy treatment classically induces cell death or cell cycle arrest. In addition to these, it is also well understood that stress signals downstream of telomere damage^5^, replication stress^6^, ribotoxic stress^7^, among others, can trigger premature reactivation of the anaphase promoting complex/cyclosome (APC/C) during G2 and allow cells to bypass mitosis and enter G1 with tetraploid (4N) DNA content. Published work shows that this M bypass largely requires p53 activation that in turn restricts continued cell cycle progression^5,8^. Therefore, M bypass largely results in permanent cell cycle arrest (i.e., senescence) when tetraploid cells are unable to re-enter S phase^6,8,9^. Since the majority of cancers have mutated p53, and these cancers tend to be harder to treat^10,11^, and in light of a recent report that demonstrated p53-independent signals are also involved in promoting M bypass^7^, we sought to understand the cell cycle fates of p53-deficient cancer cells that persist after genotoxic chemotherapy treatment.

Here, we confirm that p53-mutant cancer cells undergo widespread mitotic bypass upon exposure to genotoxic chemotherapies. Interestingly, we find that the cells that persisted after chemotherapy treatments did not undergo quiescence- or senescence-like cell cycle arrest, but instead entered an endocycle and continued to undergo successive rounds of genome duplication and mitotic bypass, ultimately becoming highly polyploid. In these models, initial M bypass allows cells to evade chemo-induced mitotic catastrophe, then continued endocycling shields DTPs from delayed cell death after drug removal. In contrast to DTPs that reside in difficult to target states of cell cycle arrest, the unique state of proliferative arrest but active cell cycle within the endocycle presents opportunity for intervention. We find that persistent DNA damage in endocycling DTPs makes them reliant on sustained G2 checkpoint signaling from WEE1 and Myt1, which promotes continued M bypass and avoidance of mitotic catastrophe. Interestingly, the roles of WEE1 and Myt1 are not redundant, and inhibiting either with a clinical-stage small molecule inhibitor is sufficient to force mitotic catastrophe and cell death in endocycling DTPs. Further, only Myt1 inhibition actively prevents M bypass, suggesting that Myt1 activity is required for premature APC/C reactivation. These results bolster previous studies in demonstrating that canonical cell cycle commitment is readily reversed under stress conditions, uncover the p53-independent drivers of this process, and define endocycling persisters as a targetable reservoir of persistence after chemotherapy treatment.

## Results

### Genotoxic chemotherapy induces widespread mitotic bypass

To monitor cell cycle dynamics in cancer cells responding to genotoxic stress, we engineered p53-mutant cancer cell lines PC3 (A138fs/del), DU145 (P223L/V274F), and MDA-MB-231 (R280K/R280K) to express the Ki67-T2A-FUCCI reporter^12^. The reporter utilizes an mCherry-Cdt1 degron and a mAG-Geminin degron to mark G1 and S/G2/M phases, respectively. The levels of each fluorescent reporter are controlled by APC/C activity, which directly degrades mAG-Geminin and indirectly leads to accumulation of mCherry-Cdt1^13^ (**Fig. 1A,B**). Expression of the reporter is driven by the Ki67 promotor and therefore non-cycling Ki67^-^ cells are identifiable as mCherry^-^/mAG^-^.

**Figure 1.**
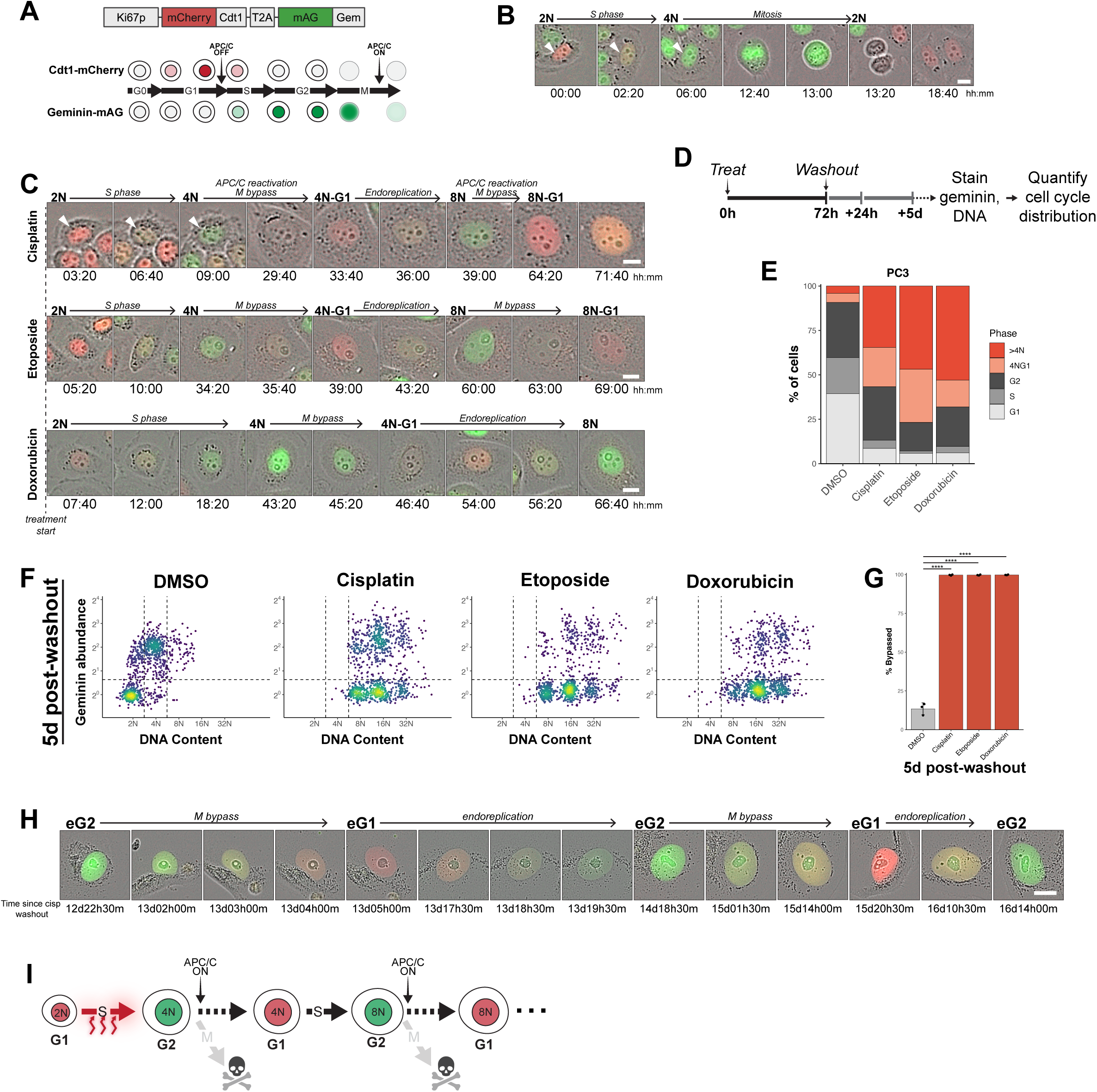
Genotoxic therapy-induced mitotic bypass precipitates persistent endocycling. **A.** Schematic of Ki67-T2A-FUCCI reporter and timeline of expression during cell cycle. **B.** Representative time series shows reporter activity during a normal cell cycle in PC3-FUCCI. Scale bar, 10μm. **C.** Representative time series shows mitotic bypass and endoreplication in PC3-FUCCI during 72h treatment with cisplatin, etoposide, or doxorubicin. Scale bar, 10μm. **D.** Experimental outline for quantifying frequency of mitotic bypass after genotoxic chemotherapy. **E.** Quantification of percent of cells in each cell cycle phase in PC3 after treatment as in (**D**). Data shown are mean percentages from 3 independent experiments. **F.** Scatter plots of DNA content vs. geminin abundance in individual surviving PC3 5d after 72h treatment with the indicated drugs. Points colored based on their relative density. Dotted lines represent cut-off values for DNA content and geminin positivity. Data shown are combined from 3 independent experiments. **G.** Quantification of the percent of cells that had bypassed mitosis in (**F**). Data are mean +/- sd from 3 independent experiments. p values calculated by one-way ANOVA and Dunnett’s post-hoc test comparing each treated condition to DMSO control. ****, p<0.0001. **H.** Representative time series of surviving PC3-FUCCI continuing to bypass mitosis and endocycle >16 days after cisplatin treatment. eG1, endocycle-G1; eG2, endocycle-G2. Scale bar, 20μm. **I.** Model of protective effect of chemotherapy-induced premature APC/C reactivation and continued endocycling. Colors of nuclei represent FUCCI reporter expression.

Timelapse imaging of FUCCI-expressing cells during treatment with LD_50_ doses of commonly used genotoxic drugs cisplatin, etoposide, and doxorubicin revealed that premature APC/C reactivation during chemo-induced G2 arrest was a widespread alternative to the expected cell cycle arrest, mitotic entry, and/or cell death (**Fig. 1C; S1A,B; and Supplementary Movie1**). These cells progress directly from G2 into tetraploid (4N) G1 without entering mitosis (no nuclear envelope breakdown or mitotic rounding), in line with previous reports of G2 cell cycle exit in response to various cellular stresses^5–7^.

To quantify the frequency of mitotic bypass in the immediate response to chemotherapy, cells were treated for 72h with cisplatin, etoposide, or doxorubicin; drug was washed out and cells that survived 24h later were fixed and stained for endogenous geminin and DAPI (**Fig. 1D**). Fluorescence intensities were extracted from individually segmented nuclei to determine the cell cycle state of each cell (**Fig. S1C**). 56.7% of cisplatin-, 76.8% of etoposide-, and 68.1% of doxorubicin-treated PC3 had bypassed mitosis, consistent with our observations in FUCCI imaging experiments (**Fig. 1E, S1D**). Similarly, the majority of MDA-MB-231 and DU145 that survived etoposide or doxorubicin treatment had bypassed mitosis, though cisplatin induced lower levels of bypass in these cells (**Fig. S1E,F**). These data demonstrate that bypass of mitosis was a frequent event in response to genotoxic chemotherapy in p53-mutant cancer cells. The prevalence of M bypass in surviving cells suggests that this is an adaptive response in the presence of genotoxic stress and is thereby selected for. Indeed, many cells that die in response to these drugs do so after failing to maintain G2 arrest and undergoing mitotic catastrophe^14^ (**Fig. S1G**). Thus, M bypass s imparts a fitness advantage during chemotherapy treatment by allowing damaged cells to circumvent mitotic cell death.

### Cells that persist after genotoxic chemotherapy enter a sustained endocycle

Cell cycle arrest, whether reversible (exit into G0, quiescence) or permanent (senescence), is a commonly reported fate of cells that survive initial chemotherapy challenge^4,15–18^. Bypassing mitosis, a reported consequence of cellular stress, can trigger permanent cell cycle exit (i.e., senescence)^9^. Here, however, despite proliferative arrest and morphological similarities to classically senescent cells, virtually all surviving cells remained Ki67^+^ and therefore continued to express the FUCCI reporter, demonstrating that cell cycle arrest was rare. Instead, most surviving cells re-entered S phase after chemo-induced M bypass. Of the PC3 that bypassed mitosis during treatment (**Fig. 1E**, red bars), 59.9% (cisplatin), 61.1% (etoposide), and 77.6% (doxorubicin), had re-entered S phase and undergone endoreduplication (second round of genome duplication without intervening mitosis), progressing past tetraploid (4N) G1 arrest (**Fig. 1E**, dark red bars). Results were similar in MDA-MB-231 and DU145 (**Fig. S1E,F**) and suggest that neither quiescence nor senescence were the primary cell fate after M bypass in surviving cells. Just 24h after drug washout there were prominent 8N G1, 16N G2, and 16N G1 populations, which require cells to have undergone a second mitotic bypass (8NG1), second endoreduplication (16NG2), and third mitotic bypass (16NG1), indicative of entry into an endocycle (**Fig. S1D**). These data indicate that although surviving cells exhibited the proliferative arrest and enlarged, flattened morphology that are hallmarks of quiescence and/or classical senescence, these were not the primary cell fates after chemotherapy challenge.

The drug doses used in these experiments result in 50% viability relative to untreated controls after 72h treatment (i.e. LD_50_), as described in the methods. In the days following drug treatment and washout, cells continue to die such that by 5 days post-drug washout, the surviving fraction is <10% of that at the end of treatment, as we have described previously^19^. At this point 5 days post-treatment an average of 99.7% (cisplatin), 99.7% (etoposide), and 99.8% (doxorubicin) of surviving PC3 had bypassed mitosis, respectively, based on analysis of DNA content and geminin abundance (**Fig. 1F,G**). On average, only 0.9%, 1.1%, and 0.8% of these cells remained in a tetraploid (4N) G1 state (i.e., mitotic bypass and subsequent arrest), confirming that endocycling was far more common than cell cycle arrest in these surviving populations. Similarly, 92.9%, 85.2%, and 74.1% of MDA-MB-231 had bypassed mitosis 5 days-post treatment with cisplatin, etoposide, or doxorubicin, respectively (**Fig. S2H**).

These persistent cells remained non-proliferative (no successful mitoses) for several weeks after drug treatment but remained Ki67^+^ and therefore continued to express the FUCCI reporter. We observed active cell cycle progression in persistent cells for the entire period of proliferative arrest after drug treatment, demonstrated by continued FUCCI reporter oscillation during time-lapse imaging of PC3-FUCCI (**Fig. 1H; Supplementary Movie2**). To confirm that FUCCI reporter oscillation accurately reported cell cycle transitions through G1-S-G2, surviving cells were pulsed with 5-ethynyl-2’-deoxyuridine (EdU) 1, 5, or 12 days after 72h cisplatin treatment and washout. Cells incorporated EdU at each timepoint, confirming progression through S phase. Moreover, discrete DNA content doublings continued over time after drug washout (**Fig. S1I**). Thus, non-proliferative persistent cells continued to undergo whole-genome replication during repeated endocycle S phases (endo-S), as the FUCCI imaging suggested.

Together, these data demonstrate that after surviving genotoxic drug treatment by bypassing mitosis and avoiding mitotic cell death, persistent cells entered a sustained endocycle: successive G1, S, and G2 phases followed by repeated M bypass (**Fig. 1I**). Endocycling persisters (DTePs) make up the predominant surviving population after treatment, demonstrating that the chemo-induced endocycle is an adaptive phenotype that is strongly selected for after genotoxic damage. We therefore asked how M bypass is controlled in these persistent cells, first during the acute response to chemotherapy and then later during sustained endocycling after drug removal.

### CDK1 inhibition is sufficient to induce premature APC/C reactivation and endocycling

Chemo-induced mitotic bypass involves 1) inhibition of G2-M progression and 2) premature APC/C reactivation during G2 arrest. Cyclin dependent kinase (CDK)1 activity drives entry into mitosis. CDK1 inhibition arrests cells in G2 and in some cases can induce premature APC/C activation and mitotic bypass^7,20–22^. We first asked whether CDK1 inhibition alone, in the absence of genotoxic stress, induced mitotic bypass and/or endocycling in our models. FUCCI-expressing cell lines were treated with the CDK1 inhibitor Ro-3306^23^ and imaged for 72h. CDK1i induced extensive APC/C reactivation during G2 (mAG-geminin degradation) and mitotic bypass in all models tested. Most of these cells proceeded to enter a subsequent S-phase (**Fig. 2A; S2A,B; and Supplementary Movie3**). Geminin immunostaining of PC3 after 72h CDK1i treatment confirmed that CDK1i alone induces widespread M bypass (>75% of cells) and endoreplication (58.2% of cells), with cells reaching ploidies ≥32N (**Fig. 2B,C**). Similar levels of endocycling occurred in MDA-MB-231 and DU145 after 72h CDK1i treatment (**Fig. S2A,B**).

**Figure 2.**
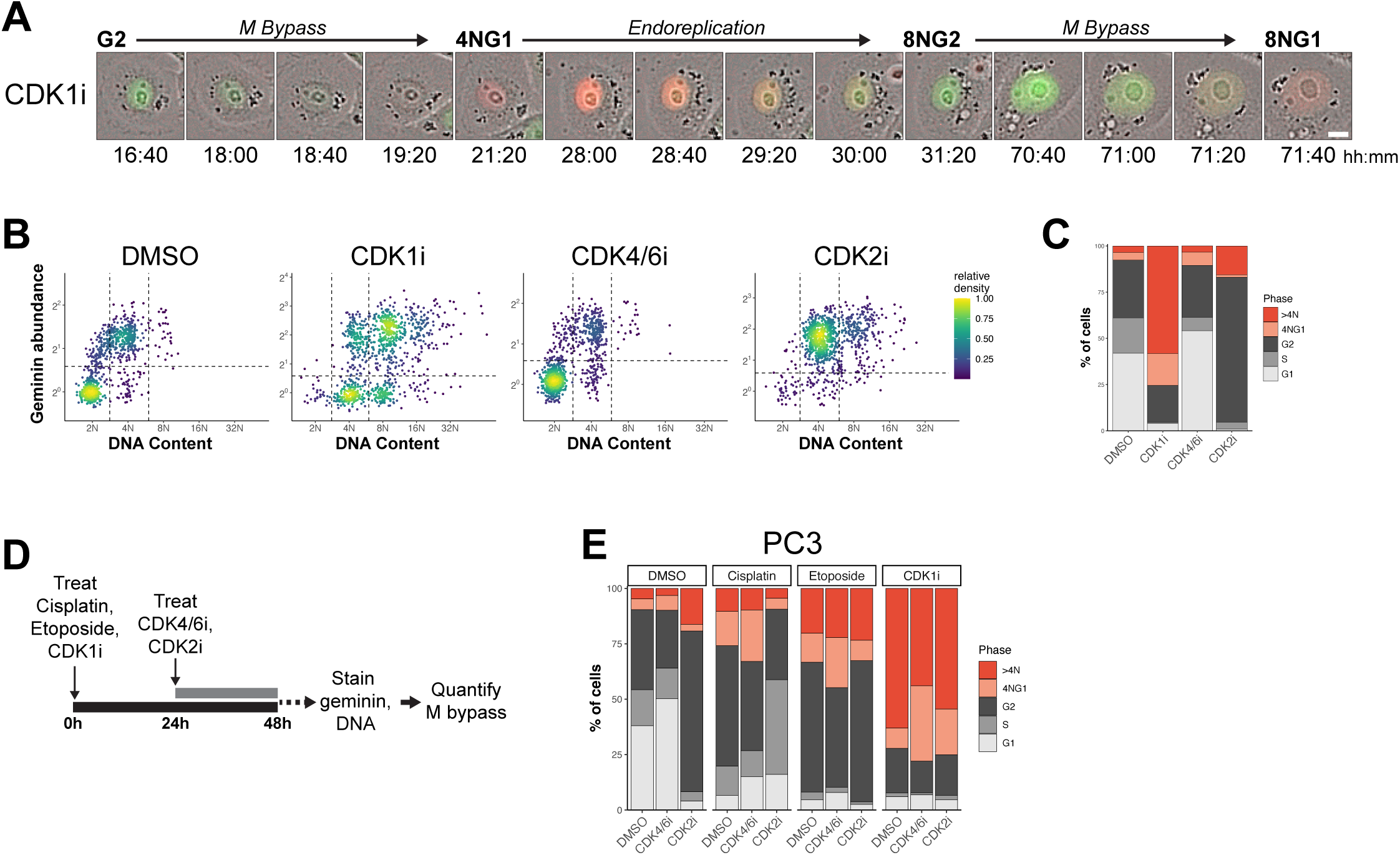
CDK1 inhibition is sufficient to induce premature APC/C reactivation and endocycling. **A.** Representative time-lapse images of PC3-FUCCI undergoing mitotic bypass and endoreplication during 72h treatment with CDK1i. Scale bar, 10μm. **B.** Scatter plots of DNA content vs. geminin abundance in individual PC3 after 72h treatment with DMSO, CDK1i, CDK4/6i, or CDK2i. Points colored based on their relative density. Dotted lines represent cut-off values for determining DNA content and geminin positivity. Data shown are combined from 3 independent experiments. 1000 individual nuclei shown for each condition. **C.** Quantification of percent of cells in each cell cycle phase in each condition from (**B**). Data shown are mean percentages from 3 independent experiments. **D.** Experimental outline for determining the effects of CDK4/6i or CDK2i on cisplatin-, etoposide-, or CDK1i-induced mitotic bypass. **E.** Percent of PC3 in each cell cycle phase after treatment as in (**D**). Data shown are mean percentages from ≥ 5 independent experiments.

Multiple recent studies have demonstrated that CDK2 and CDK4/6 activities are critical for preventing premature APC/C reactivation during G2^7,20,24^, so we next tested the roles of CDK2 and CDK4/6 activities in chemo-induced M bypass. In contrast to CDK1i, neither CDK4/6 nor CDK2 inhibition induced appreciable mitotic bypass or endoreplication in PC3 (**Fig. 2B,C**). If CDK2 and/or CDK4/6 activities prevent APC/C reactivation during G2 arrest in chemo-treated cells, then inhibiting them during chemo-induced G2 arrest would accelerate M bypass, as was shown previously in the context of other stress conditions^7^. G2 arrest was induced by 24h treatment with cisplatin, etoposide, or CDK1i (**Fig. 2B, S2C,D**) before adding CDK2i or CDK4/6i for 24h. Geminin and DNA content were analyzed to quantify the levels of M bypass in each condition (**Fig 2D**).

In PC3, CDK2i alone induced low levels of bypass in unstressed, DMSO vehicle controls and did not increase M bypass in response to cisplatin, etoposide, or CDK1i (**Fig. 2E**). Similar results were obtained in MDA-MB-231 (**Fig S2E**). CDK2, like CDK1, is a target of DNA damage signaling and we reasoned that cisplatin or etoposide treatments likely inhibit CDK2, rendering CDK2i treatment redundant. Indeed, WEE1-dependent CDK2 phosphorylation (inhibition) is induced in both cisplatin- and etoposide-treated cells (**Fig. S2F**). Since CDK2 is already inhibited by cisplatin- and etoposide-induced genotoxic stress, the redundancy of adding a CDK2 inhibitor may explain its lack of effect in those conditions, but its lack of effect in CDK1i-treated cells suggests that M bypass can proceed even when CDK2 is active. Further studies will be needed to explain how chemo-treated cells re-enter S phase after chemo-induced M bypass if CDK2 is inhibited, since CDK2 is essential for this process in other models^21^.

Inhibiting CDK4/6 during cisplatin-, etoposide-, or CDK1i-induced G2 arrest (**Fig. 2D**) only modestly increased the percent of cells that underwent M bypass in PC3 and MDA-MB-231 (**Fig. 2E; S2E**). Unlike CDK2, CDK4/6 remains active under cisplatin and etoposide treatment and continues to phosphorylate Rb in S and G2 cells regardless of their ploidy (**Fig. S2G**). Therefore, the modest effect of CDK4/6 inhibition during G2 arrest in these experiments suggests that premature APC/C reactivation occurs despite CDK4/6 activity in chemo-treated cells.

Taken together, these results demonstrate that CDK1 inhibition alone is sufficient to induce widespread M bypass and endocycling, in agreement with a previous report^21^. The modest effect of CDK4/6i on levels of M bypass in stressed cells combined with the widespread M bypass induced by CDK1i treatment alone suggests that chemo-induced CDK1 inhibition itself is the major driver of G2 arrest and premature APC/C activation in these stress conditions.

### WEE1-, Myt1- dependent CDK1 inhibition is required for drug-induced G2 arrest and M bypass

Inhibition of CDK1 is sufficient to induce M bypass and subsequent endocycling, so we sought to determine the mechanism of chemo-induced CDK1 inhibition and consequent M bypass in p53-mutant cells.

P53 and its effector p21 can induce M bypass in response to various cellular stresses by inhibiting CDK1^6,25,26^. Despite being p53-mutant, we observe an increase in p21 expression at both the mRNA and protein level in response to both cisplatin and etoposide in our models (**Fig. S3A-D**). Upregulation of p21 protein appears to be cell cycle phase-independent, with strong upregulation observed in surviving PC3 in all cell cycle phases after 72h etoposide treatment (**Fig. S3E**). Because of its reported role in mitotic bypass, we asked whether p53-independent p21 expression played a role in mitotic bypass in our models. Inducible overexpression of p21 in PC3 (PC3-p21OE) was not sufficient to induce mitotic bypass or endocycling (**Fig. S3F, G**). Further, p21 knockdown during etoposide treatment did not impair etoposide-induced M bypass or endocycling. Thus, p21 does not appear to be critical for chemo-induced M bypass or endocycling.

In the absence of functional p53, cell cycle arrest in G2 can be achieved through inhibitory phosphorylation of CDK1 at T14 by Myt1 and at Y15 by WEE1^27,28^. CDC25 phosphatases remove these marks, thus CDK1 activity is controlled by the balance of kinase and phosphatase activity, which are controlled by ataxia-telangiectasia mutated (ATM) and/or ataxia-telangiectasia and Rad3-related (ATR) ^29,30^. As expected, we observed an increase in phosphorylation of both 11hosphor-sites on CDK1 in response to cisplatin or etoposide in PC3, MDA-MB-231, and DU145, suggesting a role in mitotic bypass (**Fig. S4A-C**). After 24h cisplatin or etoposide treatment, which induced CDK1 phosphorylation and G2 arrest (**Fig. S2B,C**), inhibition of either Myt1 or WEE1 kinases was sufficient to reduce CDK1 phosphorylation on T14 or Y15, respectively. Inhibition of ATR had a similar, though more modest effect, to that of inhibiting its effector kinase WEE1 directly, while inhibiting ATM had no effect on CDK1 phosphorylation (**Fig. 3B**). Thus, site-specific CDK1 phosphorylation is dependent on Myt1i and WEE1, and inhibiting either individually is sufficient to reverse stress-induced CDK1 phosphorylation in a site-specific manner, allowing us to individually assay the functional effects of these phosphorylations on initial M bypass and subsequent endocycling.

**Figure 3.**
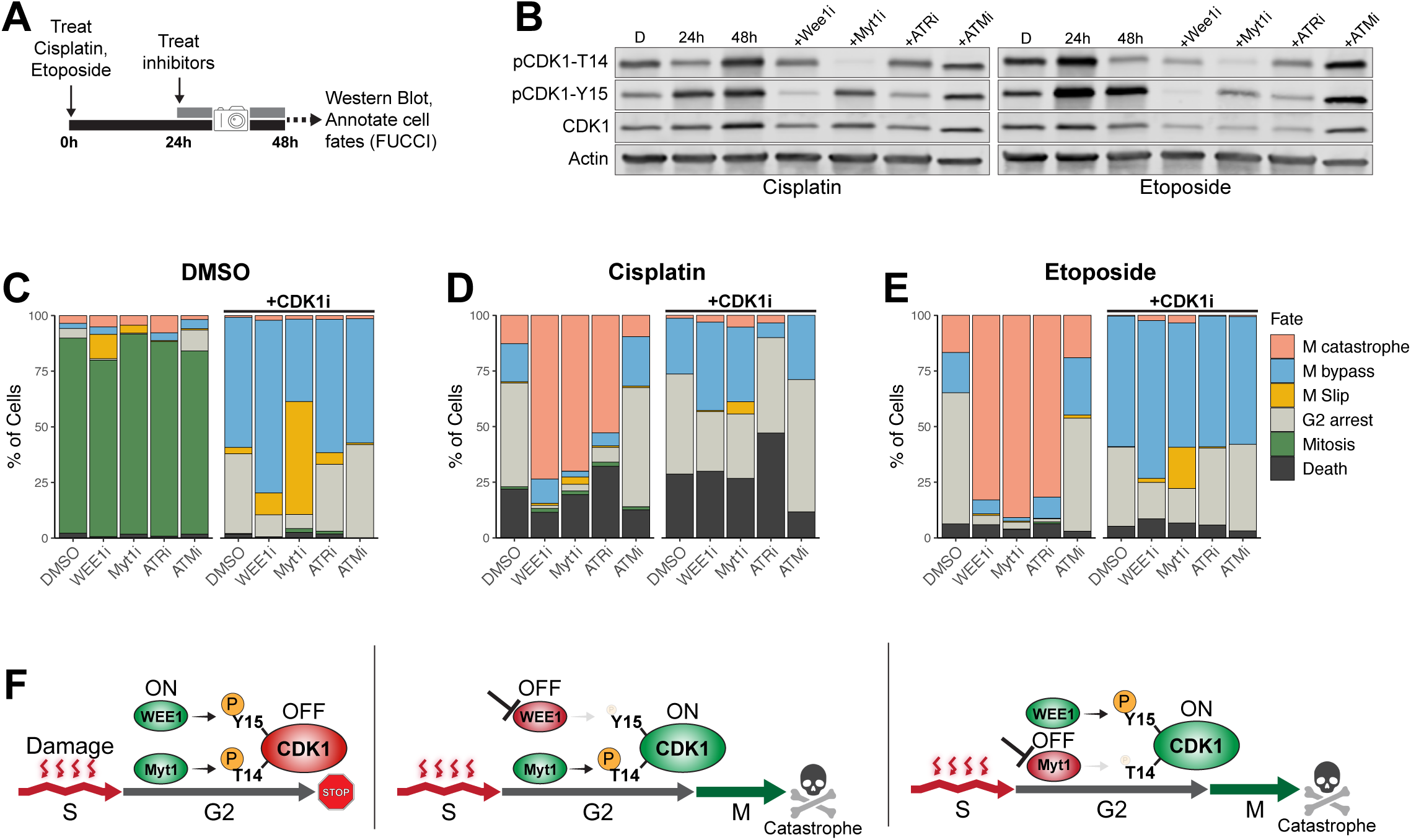
WEE1-, Myt1- dependent CDK1 inhibition is required for drug-induced G2 arrest and M bypass. **A.** Experimental outline for the analysis of the effects of WEE1, Myt1, ATR, or ATM inhibition on cisplatin or etoposide-induced CDK1 phosphorylation and cell fate. **B.** Western Blot of PC3 treated as in (**A**) showing the effects of WEE1i, Myt1i, ATRi, and ATMi on cisplatin- or etoposide-induced CDK1 phosphorylation. D, DMSO. **C-E.** Stacked bar plots showing the fates of PC3-FUCCI after DMSO (**D**), cisplatin (**E**), or etoposide (**F**) +/- inhibitor treatments as depicted in (**A**). Data shown are the mean percent of cells with each fate from ≥2 independent experiments. **F.** Model of DNA damage-induced CDK1 inhibition (left) and the effects of WEE1 or Myt1 inhibition (right).

If Myt1-dependent T14 phosphorylation or WEE1-dependent Y15 phosphorylation are critical for initial chemo-induced M bypass, then inhibition during chemo-induced G2 arrest should lead to CDK1 reactivation, forcing mitotic entry, catastrophe and cell death. Indeed, G2 checkpoint disruption is the premise that precipitated the development of both Myt1 and WEE1 inhibitors for use in combination with DNA damaging agents for cancer treatment^31–33^. Here, we sought to dissect the role of both kinases and their respective CDK1 phosphorylation sites in chemo-induced mitotic bypass and cell survival. PC3-FUCCI were treated as in (**Fig. 3A**) and time-lapse imaged for 24h following the addition of inhibitors. The resulting time series images were used to annotate cell fates. Analysis was limited to cells that were in G2 at the start of inhibitor treatment, which made up the majority of cells in agreement with geminin and DNA content data (**Fig. S2B,C**). Six distinct cell fates were observed in these experiments: successful mitoses, continued G2 arrest, cell death outside of mitosis, endo-mitosis (mitotic entry then exit before karyokinesis, resulting in a 4N-G1 cell with an intact nucleus), mitotic bypass (mAG-geminin degradation without mitotic entry), and mitotic catastrophe (either cell death during mitosis or failed karyokinesis resulting in a single G1-like cell with highly fragmented nuclei, which die within hours of mitotic failure) (**Fig. S4D**). Inhibition of Myt1, WEE1, ATR, or ATM had little effect on the fates of unstressed, DMSO-treated controls, and most cells completed successful mitoses (**Fig. 3C**). In G2 cells harboring damage from cisplatin or etoposide treatment, the addition of WEE1, Myt1, or ATR inhibitors, which disrupt CDK1 phosphorylation in a site-specific manner (**Fig. 3B**), forced widespread G2 checkpoint failure and mitotic catastrophe. ATM inhibition, which does not impair cisplatin-or etoposide-induced CDK1 phosphorylation (**Fig. 3B**), did not negatively impact G2 arrest, mitotic bypass, or survival in these cells. The addition of a CDK1 inhibitor (CDK1i) in combination with WEE1i, Myt1i, and ATRi completely reversed their effects demonstrating that the results are dependent on CDK1 activity (**Fig. 3D,E**). Although phosphorylation at either T14 or Y15 should prevent ATP binding and CDK1 activity, we found that disrupting phosphorylation at either site is sufficient to induce widespread CDK1 reactivation and mitotic catastrophe in stressed cells (**Fig. 3F**). Further, Myt1 inhibition consistently decreased cells’ ability to bypass mitosis to a greater extent than either WEE1i or ATRi (**Fig. S4E,F**), suggesting an as-yet unexplored requirement for Myt1 in stress-induced mitotic bypass. Thus, CDK1 phosphorylation-especially by Myt1-controls M bypass and thereby promotes survival in response to genotoxic drugs. The dependence of these cells on G2 checkpoint activity both highlights a well-known vulnerability in DNA-damaged cells and demonstrates that these inhibitors (especially Myt1i) also prevent stress-induced mitotic bypass. This vulnerability can be exploited to prevent M bypass and instead force extensive mitotic catastrophe, dramatically reducing the fraction of surviving cells after chemotherapy treatments, as previously described^31^.

### WEE1, Myt1 activities are required for endocycling and survival in persistent cells

CDK1 inhibition is an expected consequence of the acute stress response during chemotherapy treatment, but persistent endocycling cells continued to bypass M long after drug washout. Since we determined that WEE1 and Myt1 were critical for CDK1 inhibition, and that Myt1 was especially involved in mitotic bypass and survival in the acute response to genotoxic drugs, we asked whether these kinases similarly affected CDK1 phosphorylation and perpetuation of M bypass in endocycling persisters. In PC3 DtePs 1 or 8 days after cisplatin or etoposide treatment, inhibiting Myt1 or WEE1 strongly decreased T14 or Y15 phosphorylation, respectively, suggesting their activities continue to be required for CDK1 inhibition in endocycling persisters (**Fig. 4A; S5A**). To test the functional consequences of disrupting this site-specific CDK1 phosphorylation in endocycling persisters, PC3-FUCCI were treated with cisplatin for 3 days to induce endocycling, drug was washed out, and endocycling cells that survived 4 days later were treated with WEE1, Myt1, ATR, or ATM inhibitors and time-lapse imaged for 24h (**Fig. 4B**). Six distinct cell fates were annotated, as in (**Fig. S4D**). Disrupting either T14 or Y15 phosphorylation with Myt1i or WEE1i, respectively, was sufficient to induce widespread mitotic entry and catastrophe. ATRi had a similar effect to that of WEE1i, as expected based on its role in activating WEE1 (**Fig. 4C**). While inhibition of either WEE1, Myt1, or ATR induced widespread mitotic entry, only Myt1i reduced the fraction of cells that underwent mitotic bypass, again demonstrating a specific dependence on Myt1 activity for premature APC/C reactivation and mitotic bypass in DtePs (**Fig. 4D**). The effects of all three inhibitors were reversed by adding CDK1i, so their effects on DteP cell fate are directly related to their roles in CDK1 inhibition (**Fig. 4C**). ATM inhibition had no effect on CDK1 phosphorylation (**Fig. S5B**) or on cell fate (**Fig. S5C**) in DtePs, demonstrating ATM-independence as was seen in the acute response to drug.

**Figure 4.**
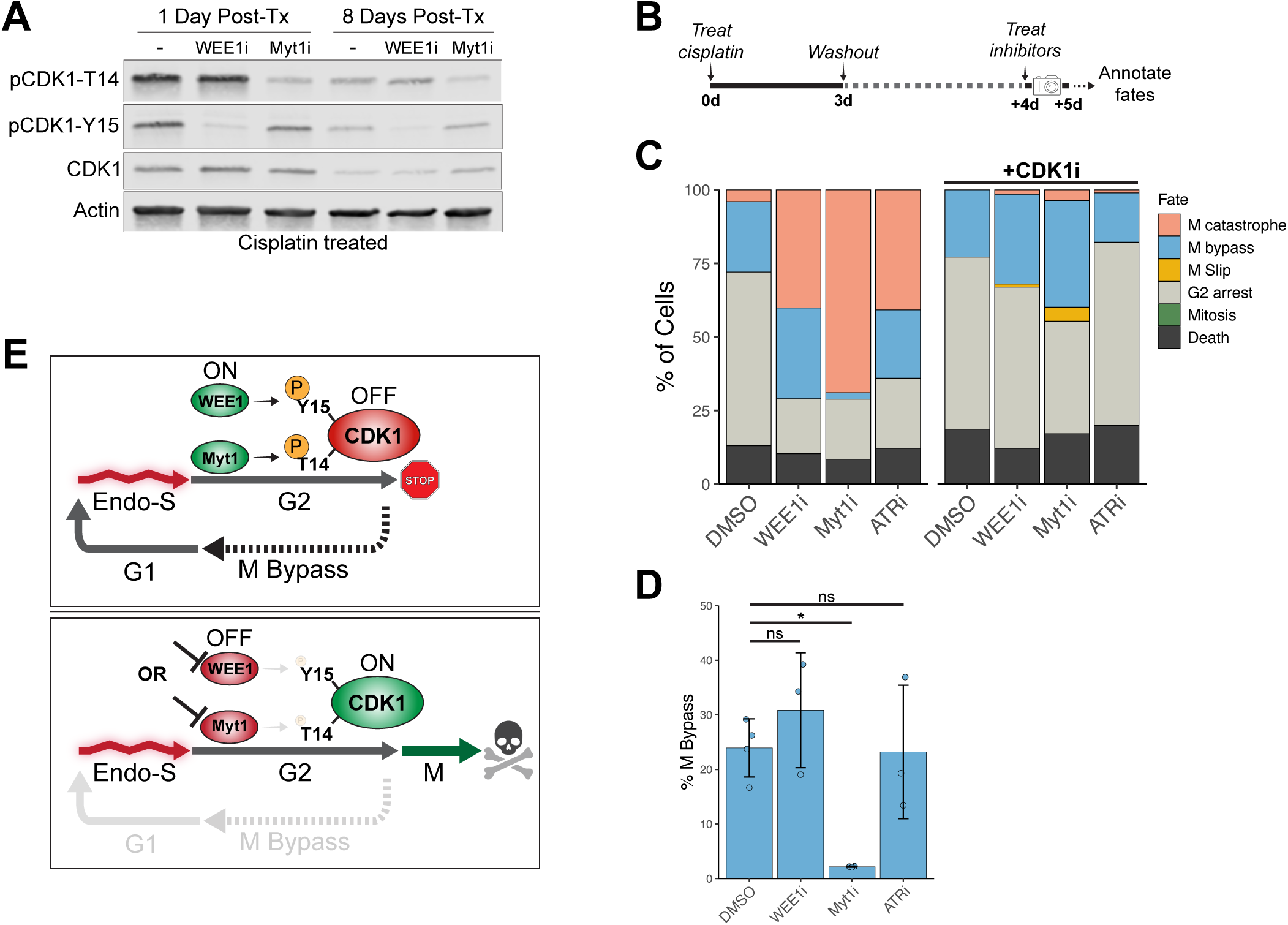
WEE1, Myt1 activities are required for endocycling and survival in persistent cells. **A.** Western Blot analysis of the effects of WEE1i or Myt1i on CDK1 phosphorylation in DTePs 1- or 8-days post cisplatin treatment. **B.** Experimental timeline for analysis of functional effects of WEE1i, Myt1i, ATRi, ATMi on DTePs: PC3-FUCCI were treated for 3 days with cisplatin then drug was washed out. 4 days later, surviving cells were treated with WEE1i, Myt1i, ATRi, or ATMi +/- CDK1i for 24 hours and timelapse imaged. Cell fates were then manually annotated. **C.** Quantification of fates of PC3-FUCCI treated as in (**B**). Data shown are the mean percent of cells with each fate from 3 individual experiments. **D.** Quantification of % M bypass from (**C**). Data are mean +/- sd from ≥3 independent experiments. p values calculated by one-way ANOVA and Dunnett’s post-hoc test comparing each treated condition to DMSO control. ns, not significant; *, p<0.05. **E.** Model of control of M bypass in DTePs.

Together, these data demonstrate that DtePs continue to have a specific dependence on Myt1- and WEE1-dependent CDK1 inhibition to enforce the G2 DNA damage checkpoint and promote their survival during the chemo-induced endocycle. Interestingly, disruption of CDK1-Y15 phosphorylation via either WEE1i or ATRi did not affect DtePs’ ability to bypass mitosis, while disruption of CDK1-T14 phosphorylation via Myt1i prevented mitotic bypass (**Fig. 4E**). These divergent responses, together with previous data suggest a previously undescribed role for Myt1 in M bypass.

### Persistent DNA damage induces stress signaling during endocycle S phases

Persistent G2 checkpoint activation after drug removal drives continued M bypass in DtePs, but it was unclear what triggers CDK1 phosphorylation well after the acute effects of genotoxic drugs. Polyploidy itself can induce DNA damage during S phase^34^, so we asked whether this was the case in DtePs. These cells do have higher levels of the DNA double-strand break (DSB) marker γH2AX and related ATM/ATR activation (phosphorylation of Chk1 and Chk2) compared to untreated cells (**Fig. 5A-F**). We classified these cells as G1 or Post-G1 based on geminin abundance to evaluate whether DNA damage and stress signaling were related to S phase entry, specifically. Levels of DNA damage were similar in G1 and Post-G1 cells, so this damage is present prior to, and is not induced as a direct consequence of endocycle S phase, in contrast to what was found in a previous report^34^ (**Fig. 5B**). Beyond their p53 mutations, there are no described defects in DNA damage repair pathways in this model, and future studies will be needed to understand the source of unresolved DNA damage in this context. In contrast, levels of p-Chk1 and p-Chk2 were both higher in Post-G1 compared to G1 cells (**Fig. 5D,F**). Therefore, we reasoned that persistent DNA damage is responsible for inducing stress signaling during endocycle S phases, downstream CDK1 inhibition, and perpetuation of M bypass in DtePs.

**Figure 5.**
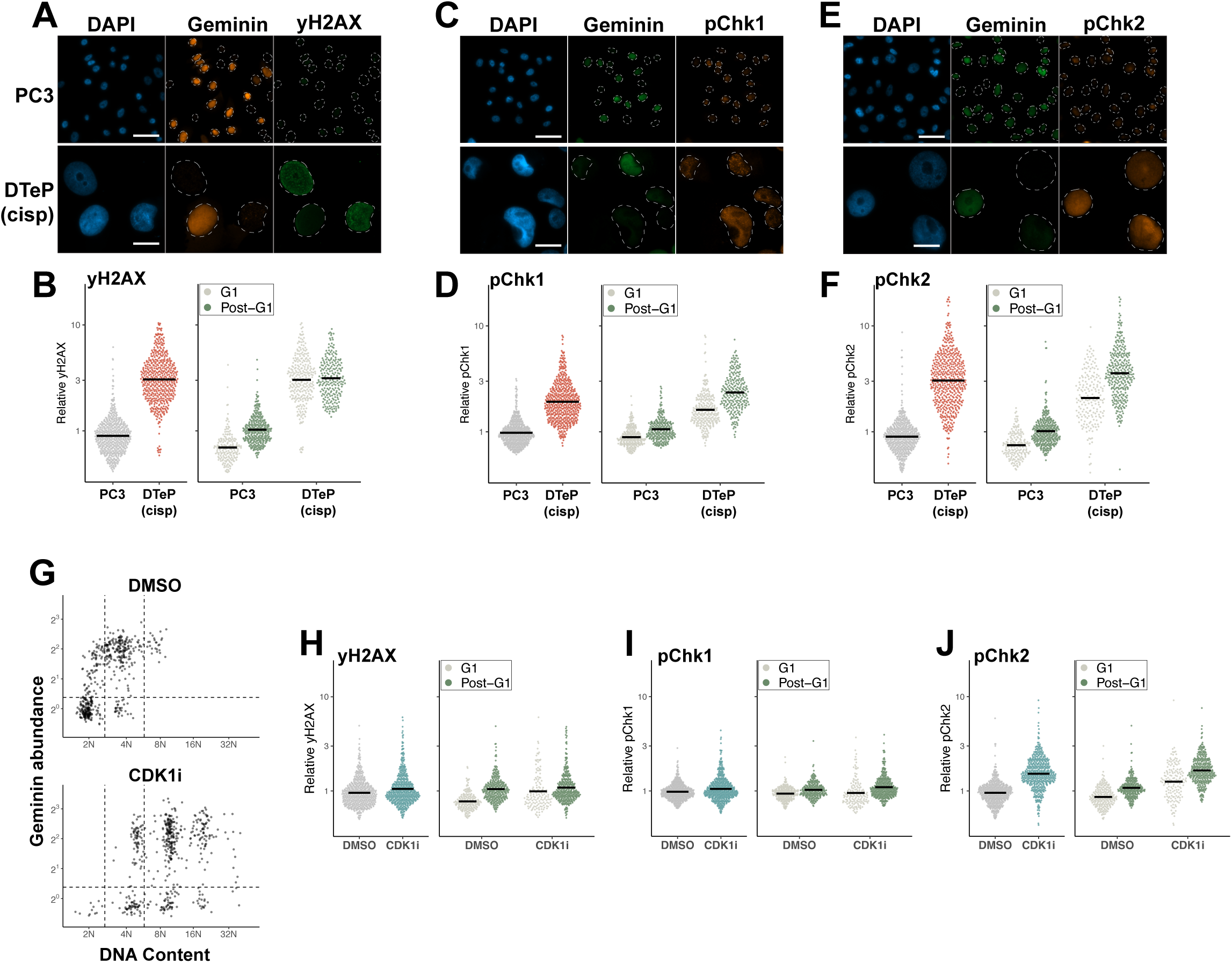
Persistent DNA damage induces stress signaling during endocycle S phases. **A.** Representative images of PC3 and DTePs 8 days post-cisplatin treatment co-immunostained for geminin and yH2AX. Scale bar, 50μm. **B.** Quantification of yH2AX abundance in individual nuclei from untreated PC3 and DTePs 8 days post-cisplatin treatment (left) and same data with cells classified as G1 or post-G1 based on geminin abundance (right). Data are combined from 2 independent experiments. Mean for each group indicated by black line. **C.** Representative images of PC3 and DTePs 8 days post-cisplatin treatment co-immunostained for geminin and pChk1. Scale bar, 50μm. **D.** Quantification of pChk1 abundance as in (**B**). **E.** Representative images of PC3 DTePs 8 days post-cisplatin treatment co-immunostained for geminin and pChk2. Scale bar, 50μm. **F.** Quantification of pChk2 abundance as in (**B**). **G.** Scatter plots of DNA content vs. geminin abundance in individual PC3 after 72h DMSO or CDK1i treatment. Data are combined from 2 independent experiments, 500 individual nuclei shown per condition. **H.** Quantification of yH2AX abundance in individual nuclei from PC3 treated 72h with DMSO or CDK1i (left) and same data with cells classified as G1 or post-G1 based on geminin abundance (right). Data are combined from 2 independent experiments, 500 individual nuclei shown per drug condition. Mean for each group indicated by black line. **I.** Quantification of pChk1 abundance in individual nuclei, as in (**H**). **J.** Quantification of pChk2 abundance in individual nuclei, as in (**H**).

We next used CDK1i-induced endocycling cells (CDK1i-ECCs) as a tool to dissect the roles of polyploidy vs. DNA damage in the activation of stress signaling during endo-S phase, as 72h CDK1i treatment induces endocycling without inducing the high levels of DNA damage seen after chemo treatments (**Fig. 5G,H**). We analyzed ATR and ATM activation, as above, in CDK1-ECCs and found these cells had only slightly elevated levels of p-Chk1 and p-Chk2, on average, and that levels were only slightly higher in post-G1 compared to G1 cells (**Fig. 5I,J**). Thus, polyploidy alone is only sufficient to induce low levels of S phase stress signaling. The modest increases in phosphorylation of Chk1 (1.01-fold increased) and Chk2 (1.63-fold increased) compared to that in cisplatin-(2.12-and 3.58-fold increases in p-Chk1 and p-Chk2, respectively) or etoposide-treated (1.97-and 2.17-fold increases in p-Chk1 and p-Chk2, respectively) endocycling persisters, suggests that the presence of persistent DNA damage likely plays the dominant role in driving endo-S phase stress signaling and persistent mitotic bypass in DtePs (**Fig. S6E**). Thus, CDK1 inhibition and endocycling are triggered by genotoxic stress, then endocycling induces further genotoxic stress and CDK1 inhibition in a circular loop that maintains a continued endocycle in persister cells

## Discussion

Chemotherapy tolerance and long-term persistence are critical drivers of cancer progression and lethality. Our results show that M bypass resulting from premature APC/C reactivation during genotoxic stress-induced G2 arrest is selected for in the presence of widely used chemotherapeutics, allowing cells to circumvent the mitotic catastrophe and cell death that these drugs induce. Stress-induced M bypass has been observed previously in other models responding to various stresses^5–7^. Our results provide further evidence that cell cycle commitment, once thought to be irreversible, is readily reversed under stressed conditions.

CDK1 inhibition induced by genotoxic stress or by a specific CDK1 inhibitor is sufficient to induce premature APC/C activation, mitotic bypass, and endocycling. CDK1 inhibition or genetic ablation is reported to induce endocycling in some, but not all models (reviewed in ^22^), together suggesting that caution should be taken in the development of CDK1 inhibition as a therapeutic strategy. Though CDK1 inhibition alone may suppress tumor growth, unintended WGD due to initiation of endocycling may exacerbate genome instability and treatment resistance. Defining biomarkers to predict which cells may be susceptible to CDK1i-induced endocycling would be an important consideration. Our results further suggest that the combination of CDK1 inhibition with genotoxic agents, as has been proposed for the treatment of metastatic pancreatic cancer^35^, for example, may be counterproductive, with CDK1 inhibition promoting M bypass and resistance to DNA damage-induced cell death.

In contrast to the well-reported states of cell cycle arrest (i.e., quiescence or senescence) that arise after chemotherapy treatment^4,15–18^, persistent cells in our model survive by accessing a sustained endocycle: a state of proliferative arrest but active cell cycle where DNA replication is uncoupled from cell division, resulting in an increasingly polyploid surviving population. Based on these results, it is likely that some previous reports of therapy-induced quiescence or senescence may include populations of endocycling cells, as has been suggested previously^36^. Future studies would benefit from employing specialized tools such as a FUCCI or other cell cycle reporters to better define “senescent” populations after drug treatment, since endocycling cells are distinct from classically G0/G1 arrested senescent cells.

Polyploid cells are thought to be refractory to the effects of DNA damage due to the protective effect of having multiple copies of each gene^37–46^. Previously published studies largely focus on tetraploid cells, and it is unknown whether protection scales with ploidy in the highly polyploid persister cells that we observe. Our data suggest that beyond polyploidy itself, the route to polyploidy (i.e., the chemo-induced endocycle) shields cells from the deleterious effects of chemotherapy and enables long-term persistence after treatment.

Critically, their reliance on sustained G2 checkpoint signaling renders endocycling persister cells vulnerable to inhibition of this checkpoint, namely by WEE1 and especially Myt1 inhibitors. This vulnerability further distinguishes endocycling persisters from cells that have undergone cell cycle arrest, as those cells reside in G1 or G0-like states and therefore have no interaction with the G2 checkpoint. This vulnerability may be clinically relevant since p53-mutated cancers that have undergone whole-genome doubling are more refractory to treatment^47^, but future studies are required to assess the relevance of endocycling DTPs *in vivo*. Inhibition of WEE1 or Myt1 separately in either the acute chemo response or in endocycling persister cells revealed that 1) disrupting phosphorylation at either of the two inhibitory sites on CDK1 was sufficient to induce CDK1 reactivation, and 2) Myt1 inhibition but not WEE1 inhibition dramatically decreases the rate of M bypass, suggesting that Myt1 activity may be required for M bypass. These differences in response may be due to dfferences in protein localization (WEE1 is nuclear, while Myt1 acts on cytoplasmic CDK1), or on Myt1’s kinase-independent role in CDK1 inhibition^48,49^, though further studies are needed define Myt1’s role in cell cycle commitment and M bypass in the context of DNA damage.

In summary, we demonstrate chemotherapy treatment commonly results in p53-independent M bypass. Since p53-deficent cells are largely incapable of senescence arrest, re-entry into S phase is common and triggers subsequent stress signaling, CDK1 inhibition, and prolonged endocycling. Cells that continue to endocycle preferentially survive and the disruption of this cycle by targeting the WEE1- and Myt1-dependent G2 checkpoint forces widespread mitotic catastrophe and cell death.

## Materials and Methods

### Cell culture

PC3 (CRL-1435), MDA-MB-231 (CRM-HTB-26), DU145 (HTB-81), and HEK293T (CRL-3216) were purchased from ATCC and cultured according to ATCC recommendations. Flasks were passaged using TrypLE Express (Gibco #12604013) when cells reached 60-80% confluence.

### Stable cell line engineering

For FUCCI lines, 1.5e6 293T were plated in 10cm dishes. The following day, cells were transfected for second generation lentivirus packaging as follows: 1ug of Hs.Ki67-T2A-FUCCI plasmid (a gift from Alexander Zambon), 1ug of psPAX2:pMD2.G (both gifts from Didier Trono, Addgene plasmid #12260, 12259) at an 8:1 ratio, and 5uL of X-tremeGENE HP transfection reagent (Roche #6366244001) were combined in 200uL in OptiMEM (Gibco 31985062) and incubated for 30 minutes before adding dropwise to 293T dishes. 18-20hr later, media was changed to target cell media + 30% heat inactivated FBS. Packaging cells were allowed to condition media for 48hr before viral harvest. Viral media was removed and filtered through a Millex HP 0.45um PES syringe filter (Millipore SLHPM33RS). Lentivirus from 1-2 10cm dishes (10-20ml) was concentrated using Clontech Lenti-X concentrator per manufacturer protocol (Takara #631232). 350k target cells were plated per well of a 6-well plate. Following concentration, viral pellet was resuspended in 2ml of target cell media + 30% heat inactivated FBS and moved to target cells in 6 well plate. 10ug/ml Polybrene (Millipore TR-1003) was added directly to well. Target cells were transduced for ∼20hr before changing media. After 3 days recovery, cells were selected with 5ug/ml blasticidin (Gibco A1113903) until untransduced control cells were dead.

For inducible p21 overexpression, Gateway donor pDONR223_CDKN1A_WT (a gift from Jesse Boehm & William Hahn & David Root; Addgene plasmid # 82201) was recombined with Gateway destination pLIX_403 (a gift from David Root; Addgene plasmid # 41395) using Gateway LR Clonase II (Invitrogen #11791020) to generate the CDKN1A expression clone.

For inducible p21 knockdown, Tet-pLKO-Neo^50^ (a gift from Dmitri Wiederschain; Addgene plasmid # 21916) [CITE] was digested with AgeI (New England Biolabs R3552) and EcoR1 (New England Biolabs R3101), then scrambled or *CDKN1A*-targeting shRNA were inserted using Instant Sticky-end Ligase Master Mix (New England Biolabs M0370) all according to manufacturer’s protocol.

Lentivirus packaging and PC3 transduction were carried out as above. 72 hours later, Tet-pLKO-Neo cells were selected with 1mg/ml G418 and pLIX403-CDKN1A cells were selected with 0.5μg/ml puromycin (pLIX403) until untransduced control cells were dead. Expression of shRNAs or CDKN1A were induced by 1μg/ml doxycycline treatment and knockdown/overexpression were confirmed by Western Blot.

<u>shRNA sequences:</u>

Scr: CCGGGCGCGATAGCGCTAATAATTTCTCGAGAAATTATTAGCGCTATCGCGCTTTTT G
sh*CDKN1A*: CCGGCGCTCTACATCTTCTGCCTTACTCGAGTAAGGCAGAAGATGTAGAGCGTTTTT G

### Drug treatments

Each cell line was treated for 72 hours with increasing doses of cisplatin (Sigma #232120), etoposide (Cell Signaling Technology #2200), or doxorubicin (Sigma #D1515) and LD50 values were determined by Alamar Blue according to manufacturer’s protocol (Invitrogen #DAL1025). The following 72h LD50 values were used for each chemotherapy drug treatment: PC3-6μM cisplatin, 5μM etoposide, 200nM doxorubicin; MDA-MB-231-12μM cisplatin, 6μM etoposide, 400nM doxorubicin; DU145-2μM cisplatin, 1.5μM etoposide, 50nM doxorubicin.

1μM WEE1i (Adavosertib, SelleckChem #S1525), 1μM Myt1i (RP-6306, MedChemExpress #HY145817A), 1μM ATRi (Ceralasertib, #S7693), 1μM ATMi (KU-60019, SelleckChem #S1570), 1μM CDK4/6i (Palbociclib, MedChemExpress #HY50767), 500nM CDK2i (INX-315, SelleckChem #E1854), and 5μM CDK1i (Ro-3306, SelleckChem #S7747) were used unless stated otherwise.

### Immunofluorescence Imaging and Cytometry analysis

Cells were cultured in 8- or 12-well chamber slides (Ibidi #80841, 81202). Cells were fixed in 4% paraformaldehyde for 15 minutes, then blocked and permeabilized with 0.3% Triton-X100 + 5% normal goat serum in PBS for 60 minutes. Primary antibody was diluted in PBS containing 1% BSA and 0.3% Triton-X100 and cells were incubated with primary antibody solution overnight at 4°C. After washing 3x 10min in PBS, fluorescent secondary antibodies along with DAPI were diluted in the same antibody buffer, cells were incubated for 60 minutes at room temperature, then washed 3x 10min in PBS before mounting with Prolong Diamond antifade mounting medium (Invitrogen P36970). Images were acquired using a Zeiss Observer Z1 microscope/ZEN pro 2.0 software (Carl Zeiss Microscopy). Cytometry analysis was performed using CellProfiler and R. Briefly, individual nuclei were segmented based on DAPI signal and intensity values within each nucleus were extracted for each fluorescent channel using CellProfiler. Downstream analysis and data visualization were performed using R.

Primary antibodies against p21 (#2947; 1:200), 23hosphor-Rb S807/11 (#8516; 1:5000), geminin (#52508; 1:1000), 23hosphor-Chk1 S345 (#2348; 1:50), and 23hosphor-Chk2 T68 (#2197; 1:100) were purchased from Cell Signaling Technologies. Anti-geminin (ab104306; 1:25000) and γH2AX (ab22551; 1:10000) were purchased from Abcam. Goat anti-mouse Alexa Fluor Plus 488 (A32723) or Alexa Fluor Plus 555 (A32727), and goat anti-rabbit Alexa Fluor Plus 488 (A32731) or Alexa Fluor Plus 555 (A32732) secondary antibodies were acquired from Invitrogen and used at 1:200.

### Immunoblotting

Cells were scraped into ice-cold PBS and pelleted by centrifugation at 5000 x g for 5 minutes. Lysates were collected in RIPA supplemented with protease/phosphatase inhibitors (Cell Signaling Technology #5872). Lysates were clarified by centrifugation at 10000 x g for 15 minutes at 4°C. Protein concentration was measured using Pierce BCA assay (Thermo Scientific #23225) according to manufacturer protocol. Normalized samples were mixed with 4x Laemmli buffer (BioRad #1610747) and boiled at 95°C for 10 minutes before being loaded into 4-20% Bis-tris polyacrylamide gels (BioRad #4561095). Electrophoresis performed with Tris/Glycine/SDS buffer (BioRad #1610732). Protein was transferred to nitrocellulose membranes (BioRad #1704158) using the BioRad TransBlot Turbo system per manufacturer protocol. Membranes were blocked for 60 minutes in StartingBlock buffer (Thermo Scientific #37538). Primary antibody was diluted in Intercept T20 buffer (Licor #927-65001) and membranes were incubated overnight at 4°C with agitation followed by 3x 5min washes with TBST, then incubated 60 minutes in secondary antibody at room temperature, washed 3x 5min with TBST, and imaged with Licor Odyssey imager. Primary antibodies against 24hosphor-CDK1 Y15 (#4539), CDK1 (#9116), p21 (#2947), and CDK2 (#18048) were purchased from Cell Signaling Technology and used at 1:1000. Anti-24hosphor-CDK T14 (#ab58509) and 24hosphor-CDK2 Y15 (ab314034) were purchased from Abcam and used at 1:1000. Anti-actin (#A5441) was purchased from Sigma and used at 1:5000. Goat anti-Rabbit 800CW (#926-32211) and goat anti-mouse 680RD (#926-68070) secondary antibodies were purchased from Licor and used at 1:20000.

### Time lapse imaging

Cells were plated in 96-well ImageLock plates (Sartorius #BA-04856) in phenol-red free medium and imaged every 20 minutes with Incucyte SX5 equipped with Green/Orange/NIR optical module (Sartorius).

### qPCR

Cells were rinsed with ice-cold PBS then scraped into 1mL ice-cold PBS and pelleted by centrifugation at 1000 x g for 5 minutes at 4°C. mRNA was isolated using Rneasy Plus kit (Qiagen #74134) and concentration was quantified with NanoDrop One (Thermo Scientific). Reverse transcription of 1ug input mRNA was done using iScript cDNA synthesis kit (BioRad #1708890) according to manufacturer’s protocol. Real-time PCR was done using SssoFast EvaGreen (BioRad #1725200) and run on BioRad CFX Opus real-time PCR machine. The following primers were purchased from Integrated DNA technologies: CDKN1A (fwd: CGGAACAAGGAGTCAGACATT; rev: AGTGCCAGGAAAGACAACTAC), ACTB (fwd: CATGTACGTTGCTATCCAGGC; rev: CTCCTTAATGTCACGCACGAT).

### EdU incorporation assay

Cells were plated in 8-well chamber slides (Ibidi #80841) and 24h later pulsed with 10uM EdU (Molecular Probes E10187) for 20h. Click-It Plus EdU assay (Molecular Probes C10640) was performed according to manufacturer’s protocol then samples were incubated with DAPI for 10 minutes before mounting slides with Prolong Diamond antifade mounting medium (Invitrogen P36970). Images were acquired and analyzed as for Immunofluorescence experiments, above.

### Statistical analysis

All statistical analyses were performed in R with α = 0.05. Specific comparisons, tests used, and p values are described in each figure legend.

## Author contributions

KT, KJP, and SRA designed the study; KT, LR, and MEY performed the experiments; KT, LR, and GB analyzed the data, and KT wrote the manuscript with input from KJP and SRA; KJP and SRA acquired funding.

## Acknowledgements

We would like to thank Dr. Sergi Regot and Dr. Christina Ferrer for helpful discussion and the current and former members of the Pienta and Amend groups for critical feedback. This work was funded by US Department of Defense CDMRP/PCRP (W81XWH-20-10353 and W81XWH-22-1-0680), the Prostate Cancer Foundation, and the Patrick C. Walsh Prostate Cancer Research Fund to SRA; and NIH/NCI P01CA093900 and the Prostate Cancer Foundation to KJP.

## Competing interests

The authors declare no competing interests.

## Ethics

All experiments were carried out using established human cell lines obtained from ATCC. Cells lines were routinely tested for mycoplasma and identities were authenticated using STR typing.

## Notes

### Competing Interest Statement

The authors have declared no competing interest.

### Summary of Updates

Figures and text revised for clarity; supplemental information added to provide more context

## References

1. Marine, J.C., Dawson, S.J., and Dawson, M.A. (2020). Non-genetic mechanisms of therapeutic resistance in cancer. Nat Rev Cancer 20, 743–756. 10.1038/s41568-020-00302-4.

2. Pu, Y., Li, L., Peng, H., Liu, L., Heymann, D., Robert, C., Vallette, F., and Shen, S. (2023). Drug-tolerant persister cells in cancer: the cutting edges and future directions. Nat Rev Clin Oncol 20, 799–813. 10.1038/s41571-023-00815-5.

3. Shen, S., Vagner, S., and Robert, C. (2020). Persistent Cancer Cells: The Deadly Survivors. Cell 183, 860–874. 10.1016/j.cell.2020.10.027.

4. Russo, M., Chen, M., Mariella, E., Peng, H., Rehman, S.K., Sancho, E., Sogari, A., Toh, T.S., Balaban, N.Q., Batlle, E., et al. (2024). Cancer drug-tolerant persister cells: from biological questions to clinical opportunities. Nat Rev Cancer 24, 694–717. 10.1038/s41568-024-00737-z.

5. Davoli, T., Denchi, E.L., and de Lange, T. (2010). Persistent telomere damage induces bypass of mitosis and tetraploidy. Cell 141, 81–93. 10.1016/j.cell.2010.01.031.

6. Zeng, J., Hills, S.A., Ozono, E., and Diffley, J.F.X. (2023). Cyclin E-induced replicative stress drives p53-dependent whole-genome duplication. Cell 186, 528–542 e514. 10.1016/j.cell.2022.12.036.

7. McKenney, C., Lendner, Y., Guerrero Zuniga, A., Sinha, N., Veresko, B., Aikin, T.J., and Regot, S. (2024). CDK4/6 activity is required during G(2) arrest to prevent stress-induced endoreplication. Science 384, eadi2421. 10.1126/science.adi2421.

8. Ganem, N.J., and Pellman, D. (2007). Limiting the proliferation of polyploid cells. Cell 131, 437–440. 10.1016/j.cell.2007.10.024.

9. Johmura, Y., Shimada, M., Misaki, T., Naiki-Ito, A., Miyoshi, H., Motoyama, N., Ohtani, N., Hara, E., Nakamura, M., Morita, A., et al. (2014). Necessary and sufficient role for a mitosis skip in senescence induction. Mol Cell 55, 73–84. 10.1016/j.molcel.2014.05.003.

10. Zhou, X., Hao, Q., and Lu, H. (2019). Mutant p53 in cancer therapy-the barrier or the path. J Mol Cell Biol 11, 293–305. 10.1093/jmcb/mjy072.

11. Hientz, K., Mohr, A., Bhakta-Guha, D., and Efferth, T. (2017). The role of p53 in cancer drug resistance and targeted chemotherapy. Oncotarget 8, 8921–8946. 10.18632/oncotarget.13475.

12. Zambon, A.C., Hsu, T., Kim, S.E., Klinck, M., Stowe, J., Henderson, L.M., Singer, D., Patam, L., Lim, C., McCulloch, A.D., et al. (2020). Methods and sensors for functional genomic studies of cell-cycle transitions in single cells. Physiol Genomics 52, 468–477. 10.1152/physiolgenomics.00065.2020.

13. Peters, J.M. (2006). The anaphase promoting complex/cyclosome: a machine designed to destroy. Nat Rev Mol Cell Biol 7, 644–656. 10.1038/nrm1988.

14. Vakifahmetoglu, H., Olsson, M., and Zhivotovsky, B. (2008). Death through a tragedy: mitotic catastrophe. Cell Death Differ 15, 1153–1162. 10.1038/cdd.2008.47.

15. Guillon, J., Petit, C., Toutain, B., Guette, C., Lelievre, E., and Coqueret, O. (2019). Chemotherapy-induced senescence, an adaptive mechanism driving resistance and tumor heterogeneity. Cell Cycle 18, 2385–2397. 10.1080/15384101.2019.1652047.

16. Duy, C., Li, M., Teater, M., Meydan, C., Garrett-Bakelman, F.E., Lee, T.C., Chin, C.R., Durmaz, C., Kawabata, K.C., Dhimolea, E., et al. (2021). Chemotherapy Induces Senescence-Like Resilient Cells Capable of Initiating AML Recurrence. Cancer Discov 11, 1542–1561. 10.1158/2159-8290.CD-20-1375.

17. Ashraf, H.M., Fernandez, B., and Spencer, S.L. (2023). The intensities of canonical senescence biomarkers integrate the duration of cell-cycle withdrawal. Nat Commun 14, 4527. 10.1038/s41467-023-40132-0.

18. Fernandez, B., Passanisi, V.J., Ashraf, H.M., and Spencer, S.L. (2025). Single-cell RNA sequencing reveals a quiescence-senescence continuum and distinct senotypes following chemotherapy. Nat Commun 17, 169. 10.1038/s41467-025-66836-z.

19. Kim, C.J., Gonye, A.L., Truskowski, K., Lee, C.F., Cho, Y.K., Austin, R.H., Pienta, K.J., and Amend, S.R. (2023). Nuclear morphology predicts cell survival to cisplatin chemotherapy. Neoplasia 42, 100906. 10.1016/j.neo.2023.100906.

20. Cornwell, J.A., Crncec, A., Afifi, M.M., Tang, K., Amin, R., and Cappell, S.D. (2023). Loss of CDK4/6 activity in S/G2 phase leads to cell cycle reversal. Nature 619, 363–370. 10.1038/s41586-023-06274-3.

21. Ma, H.T., Tsang, Y.H., Marxer, M., and Poon, R.Y. (2009). Cyclin A2-cyclin-dependent kinase 2 cooperates with the PLK1-SCFbeta-TrCP1-EMI1-anaphase-promoting complex/cyclosome axis to promote genome reduplication in the absence of mitosis. Mol Cell Biol 29, 6500–6514. 10.1128/MCB.00669-09.

22. Porter, A.C. (2008). Preventing DNA over-replication: a Cdk perspective. Cell Div 3, 3. 10.1186/1747-1028-3-3.

23. Vassilev, L.T., Tovar, C., Chen, S., Knezevic, D., Zhao, X., Sun, H., Heimbrook, D.C., and Chen, L. (2006). Selective small-molecule inhibitor reveals critical mitotic functions of human CDK1. Proc Natl Acad Sci U S A 103, 10660–10665. 10.1073/pnas.0600447103.

24. Kim, K., Armand, J., Kim, S., and Yang, H.W. (2025). E2F activity determines mitosis versus whole-genome duplication in G2-arrested cells. Nat Commun 16, 6677. 10.1038/s41467-025-62061-w.

25. Bates, S., Ryan, K.M., Phillips, A.C., and Vousden, K.H. (1998). Cell cycle arrest and DNA endoreduplication following p21Waf1/Cip1 expression. Oncogene 17, 1691–1703. 10.1038/sj.onc.1202104.

26. Ullah, Z., Lee, C.Y., and Depamphilis, M.L. (2009). Cip/Kip cyclin-dependent protein kinase inhibitors and the road to polyploidy. Cell Div 4, 10. 10.1186/1747-1028-4-10.

27. Liu, F., Stanton, J.J., Wu, Z., and Piwnica-Worms, H. (1997). The human Myt1 kinase preferentially phosphorylates Cdc2 on threonine 14 and localizes to the endoplasmic reticulum and Golgi complex. Mol Cell Biol 17, 571–583. 10.1128/MCB.17.2.571.

28. Parker, L.L., Atherton-Fessler, S., and Piwnica-Worms, H. (1992). p107wee1 is a dual-specificity kinase that phosphorylates p34cdc2 on tyrosine 15. Proc Natl Acad Sci U S A 89, 2917–2921. 10.1073/pnas.89.7.2917.

29. Boutros, R., Dozier, C., and Ducommun, B. (2006). The when and wheres of CDC25 phosphatases. Curr Opin Cell Biol 18, 185–191. 10.1016/j.ceb.2006.02.003.

30. Blackford, A.N., and Jackson, S.P. (2017). ATM, ATR, and DNA-PK: The Trinity at the Heart of the DNA Damage Response. Mol Cell 66, 801–817. 10.1016/j.molcel.2017.05.015.

31. Aarts, M., Sharpe, R., Garcia-Murillas, I., Gevensleben, H., Hurd, M.S., Shumway, S.D., Toniatti, C., Ashworth, A., and Turner, N.C. (2012). Forced mitotic entry of S-phase cells as a therapeutic strategy induced by inhibition of WEE1. Cancer Discov 2, 524–539. 10.1158/2159-8290.CD-11-0320.

32. Hirai, H., Arai, T., Okada, M., Nishibata, T., Kobayashi, M., Sakai, N., Imagaki, K., Ohtani, J., Sakai, T., Yoshizumi, T., et al. (2010). MK-1775, a small molecule Wee1 inhibitor, enhances anti-tumor efficacy of various DNA-damaging agents, including 5-fluorouracil. Cancer Biol Ther 9, 514–522. 10.4161/cbt.9.7.11115.

33. Szychowski, J., Papp, R., Dietrich, E., Liu, B., Vallee, F., Leclaire, M.E., Fourtounis, J., Martino, G., Perryman, A.L., Pau, V., et al. (2022). Discovery of an Orally Bioavailable and Selective PKMYT1 Inhibitor, RP-6306. J Med Chem 65, 10251–10284. 10.1021/acs.jmedchem.2c00552.

34. Gemble, S., Wardenaar, R., Keuper, K., Srivastava, N., Nano, M., Mace, A.S., Tijhuis, A.E., Bernhard, S.V., Spierings, D.C.J., Simon, A., et al. (2022). Genetic instability from a single S phase after whole-genome duplication. Nature 604, 146–151. 10.1038/s41586-022-04578-4.

35. Wang, Q., Bode, A.M., and Zhang, T. (2023). Targeting CDK1 in cancer: mechanisms and implications. NPJ Precis Oncol 7, 58. 10.1038/s41698-023-00407-7.

36. Hayashi, K., Horisaka, K., Harada, Y., Ogawa, Y., Yamashita, T., Kitano, T., Wakita, M., Fukusumi, T., Inohara, H., Hara, E., and Matsumoto, T. (2024). Polyploidy mitigates the impact of DNA damage while simultaneously bearing its burden. Cell Death Discov 10, 436. 10.1038/s41420-024-02206-w.

37. Darmasaputra, G.S., van Rijnberk, L.M., and Galli, M. (2024). Functional consequences of somatic polyploidy in development. Development 151. 10.1242/dev.202392.

38. Dewhurst, S.M., McGranahan, N., Burrell, R.A., Rowan, A.J., Gronroos, E., Endesfelder, D., Joshi, T., Mouradov, D., Gibbs, P., Ward, R.L., et al. (2014). Tolerance of whole-genome doubling propagates chromosomal instability and accelerates cancer genome evolution. Cancer Discov 4, 175–185. 10.1158/2159-8290.CD-13-0285.

39. Hassel, C., Zhang, B., Dixon, M., and Calvi, B.R. (2014). Induction of endocycles represses apoptosis independently of differentiation and predisposes cells to genome instability. Development 141, 112–123. 10.1242/dev.098871.

40. Kuznetsova, A.Y., Seget, K., Moeller, G.K., de Pagter, M.S., de Roos, J.A., Durrbaum, M., Kuffer, C., Muller, S., Zaman, G.J., Kloosterman, W.P., and Storchova, Z. (2015). Chromosomal instability, tolerance of mitotic errors and multidrug resistance are promoted by tetraploidization in human cells. Cell Cycle 14, 2810–2820. 10.1080/15384101.2015.1068482.

41. Lopez, S., Lim, E.L., Horswell, S., Haase, K., Huebner, A., Dietzen, M., Mourikis, T.P., Watkins, T.B.K., Rowan, A., Dewhurst, S.M., et al. (2020). Interplay between whole-genome doubling and the accumulation of deleterious alterations in cancer evolution. Nat Genet 52, 283–293. 10.1038/s41588-020-0584-7.

42. Schmidt, M.J., Naghdloo, A., Prabakar, R.K., Kamal, M., Cadaneanu, R., Garraway, I.P., Lewis, M., Aparicio, A., Zurita-Saavedra, A., Corn, P., et al. (2025). Polyploid cancer cells reveal signatures of chemotherapy resistance. Oncogene 44, 439–449. 10.1038/s41388-024-03212-z.

43. Herriage, H.C., Huang, Y.T., and Calvi, B.R. (2024). The antagonistic relationship between apoptosis and polyploidy in development and cancer. Semin Cell Dev Biol 156, 35–43. 10.1016/j.semcdb.2023.05.009.

44. Mehrotra, S., Maqbool, S.B., Kolpakas, A., Murnen, K., and Calvi, B.R. (2008). Endocycling cells do not apoptose in response to DNA rereplication genotoxic stress. Genes Dev 22, 3158–3171. 10.1101/gad.1710208.

45. Zheng, L., Dai, H., Zhou, M., Li, X., Liu, C., Guo, Z., Wu, X., Wu, J., Wang, C., Zhong, J., et al. (2012). Polyploid cells rewire DNA damage response networks to overcome replication stress-induced barriers for tumour progression. Nat Commun 3, 815. 10.1038/ncomms1825.

46. Shu, Z., Row, S., and Deng, W.M. (2018). Endoreplication: The Good, the Bad, and the Ugly. Trends Cell Biol 28, 465–474. 10.1016/j.tcb.2018.02.006.

47. Hobor, S., Al Bakir, M., Hiley, C.T., Skrzypski, M., Frankell, A.M., Bakker, B., Watkins, T.B.K., Markovets, A., Dry, J.R., Brown, A.P., et al. (2024). Mixed responses to targeted therapy driven by chromosomal instability through p53 dysfunction and genome doubling. Nat Commun 15, 4871. 10.1038/s41467-024-47606-9.

48. Liu, F., Rothblum-Oviatt, C., Ryan, C.E., and Piwnica-Worms, H. (1999). Overproduction of human Myt1 kinase induces a G2 cell cycle delay by interfering with the intracellular trafficking of Cdc2-cyclin B1 complexes. Mol Cell Biol 19, 5113–5123. 10.1128/MCB.19.7.5113.

49. Wells, N.J., Watanabe, N., Tokusumi, T., Jiang, W., Verdecia, M.A., and Hunter, T. (1999). The C-terminal domain of the Cdc2 inhibitory kinase Myt1 interacts with Cdc2 complexes and is required for inhibition of G(2)/M progression. J Cell Sci 112 ( *Pt 19*), 3361–3371. 10.1242/jcs.112.19.3361.

50. Wiederschain, D., Wee, S., Chen, L., Loo, A., Yang, G., Huang, A., Chen, Y., Caponigro, G., Yao, Y.M., Lengauer, C., et al. (2009). Single-vector inducible lentiviral RNAi system for oncology target validation. Cell Cycle 8, 498–504. 10.4161/cc.8.3.7701.

