## Supplemental Figures for "Mitotic bypass and continued endocycling promote cancer cell survival after genotoxic chemotherapy"

**Figure S1**

**A.** Representative time series shows mitotic bypass and endoreplication in MDA-MB-231-FUCCI during 72h treatment with cisplatin, etoposide, or doxorubicin. Scale bar, 10μm.

**B.** Representative time series shows mitotic bypass and endoreplication in DU145-FUCCI during 72h treatment with cisplatin, etoposide, or doxorubicin. Scale bar, 10μm.

**C.** Model of DNA content vs. geminin abundance scatterplot with gating strategy and interpretation: 2N, geminin$^-$ cells are in G1; between 2N and 4N, geminin$^+$ cells are in S; 4N, geminin$^+$ cells are in G2; 4N, geminin$^-$ cells have bypassed M into 4N-G1; cells with >4N DNA content have endoreduplicated.

**D.** Scatter plots of DNA content vs. geminin abundance in individual surviving PC3 24h after 72h treatment with the indicated drugs. Points colored based on their relative density. Dotted lines represent cut-off values for DNA content and geminin positivity. Data shown are combined from 3 independent experiments, 1000 individual data points shown for each condition.

**E.** Quantification of the percent of MDA-MB-231 in each cell cycle phase after treatment as in **Fig. 1D**. Data shown are mean percentages from 3 independent experiments.

**F.** Quantification of the percent of DU145 in each cell cycle phase after treatment as in **Fig. 1D**. Data shown are mean percentages from 3 independent experiments.

**G.** Representative time series of PC3-FUCCI undergoing mitotic catastrophe in response to cisplatin treatment. Scale bar, 10μm.

**H.** Quantification of the percent of MDA-MB-231 that had bypassed mitosis 5d after 72h treatment with the indicated drugs. Data are mean +/- sd from ≥2 independent experiments. p values calculated by one-way ANOVA and Dunnett's post-hoc test comparing each treated condition to DMSO control. ***, $p<0.001$; ****, $p<0.0001$.

**I.** Representative images of DAPI and EdU incorporation and DNA content quantification in surviving PC3 1, 5, or 12 days after cisplatin treatment. Scale bar, 50μm.

**Figure S2**

**A.** Representative time-lapse images of MDA-MB-231 undergoing M bypass and endoreplication during 72h treatment with CDK1i and quantification of percent of cells in each cell cycle phase after 72h DMSO or CDK1i treatment. Scale bar, 10μm. Data shown are mean percentages from 3 independent experiments.

**B.** Representative time-lapse images of DU145 undergoing M bypass and endoreplication during 72h treatment with CDK1i and quantification of percent of cells in each cell cycle phase after 72h DMSO or CDK1i treatment. Scale bar, 10μm. Data shown are mean percentages from 3 independent experiments.

**C.** Scatter plots of DNA content vs. geminin abundance in PC3 after 24h treatment with DMSO, CDK1i, cisplatin, or etoposide. Points colored based on their relative density. Dotted lines represent cut-off values for determining DNA content and geminin positivity. 1000 individual nuclei shown for each condition.

**D.** Quantification of percent of cells in each phase from (**C**).

**E.** Percent of MDA-MB-231 in each cell cycle phase after treatment as in (**Fig. 2D**). Data shown are mean percentages from 2 independent experiments.

**F.** Western Blot analysis of WEE1-dependent CDK2 phosphorylation in (left) cisplatin- or (right) etoposide- treated PC3

**G.** Scatter plots of DNA content vs. geminin abundance in PC3 after 72h treatment with DMSO, cisplatin, etoposide, or CDK1i. Points colored based on p-Rb abundance.

**Figure S3**

**A.** Western Blot of p21 expression in PC3 after 24-, 48-, or 72-hour treatment with cisplatin or etoposide.

**B.** Relative expression of *CDKN1A* mRNA in PC3 after 72h treatment with cisplatin or etoposide compared to DMSO-treated control.

**C.** Western Blot of p21 expression in MDA-MB-231 after 24-, 48-, or 72-hour treatment with cisplatin or etoposide

**D.** Relative expression of *CDKN1A* mRNA in MDA-MB-231 after 72h treatment with cisplatin or etoposide compared to DMSO-treated control.

**E.** PC3 treated 72h with DMSO or etoposide, stained with DAPI, and co-immunostained for geminin and p21, to determine cell cycle phase and p21 abundance in each nucleus. Data shown are p21 abundances in individual PC3 in each cell cycle phase after treatment.

**F.** Western Blot of p21 expression in PC3-WT and PC3-p21OE after 72h vehicle or doxycycline treatment to induce shRNA expression.

**G.** Quantification of M bypassed cells / total cells in PC3-WT and PC3-p21OE after 72h vehicle or doxycycline treatment to induce shRNA expression. Data are mean +/- sd from 3 individual experiments. p values calculated by unpaired t test comparing - / + doxycycline conditions in each group. ns, not significant.

**H.** Western Blot of p21 expression in PC3-shScr and PC3-shCDKN1A after 72hr treatment with DMSO or etoposide +/- doxycycline to induce p21 overexpression.

**I.** Quantification of M bypassed cells / total cells in PC3-shScr and PC3-shCDKN1A after 72hr treatment with DMSO or etoposide +/- doxycycline. Data are mean +/- sd from 3 individual experiments. p values calculated by unpaired t test comparing - /+ doxycycline conditions within each group. ns, not significant.

**Figure S4**

**A.** Western Blot analysis of CDK1 phosphorylation in PC3 after 24, 48, or 72hr treatment with cisplatin or etoposide. D, DMSO; p, phosphorylation.

**B.** Western Blot analysis of CDK1 phosphorylation in MDA-MB-231 after 24, 48, or 72hr treatment with cisplatin or etoposide. D, DMSO; p, phosphorylation.

**C.** Western Blot analysis of CDK1 phosphorylation in DU145 after 24, 48, or 72hr treatment with cisplatin or etoposide. D, DMSO; p, phosphorylation.

**D.** Example time-lapse images illustrating PC3-FUCCI cell fates during treatment as in **Fig. 3A**.

**E.** Quantification of % M bypass from **Fig. 3D**. Data are mean +/- sd from ≥2 independent experiments. p values calculated by one-way ANOVA and Dunnett's post-hoc test comparing each treated condition to DMSO control. ns, not significant; **, p<0.01.

**F.** Quantification of % M bypass from **Fig. 3E**. Data are mean +/- sd from ≥2 independent experiments. p values calculated by one-way ANOVA and Dunnett's post-hoc test comparing each treated condition to DMSO control. ns, not significant; *, p<0.05; **, p<0.01; ***, p<0.001.

**Figure S5**

**A.** Western Blot analysis of the effects of inhibiting WEE1 or Myt1 on CDK1 phosphorylation in DTePs 1- or 8-days post etoposide treatment.

**B.** Western Blot analysis of the effects of ATMi on CDK1 phosphorylation in DTePs 4 days post-cisplatin or etoposide treatment.

**C.** Cell fate analysis of cisplatin-treated DTePs after ATMi +/- CDK1i treatment as in (**Fig. 4B**).

**Figure S6**

**A.** Example quantification of geminin abundance in PC3 and DTePs 1 or 8 days post treatment to demonstrate cell cycle classification of G1 or Post-G1 based on geminin abundance.

**B.** Quantification of yH2AX abundance in individual nuclei from untreated PC3 and DTePs 8 days post-etoposide treatment (left) and same data with cells classified as G1 or post-G1 based on geminin abundance (right). Data are combined from 2 independent experiments. Mean for each group indicated by black line.

**C.** Quantification of p-Chk1 abundance in individual nuclei as in (**B**).

**D.** Quantification of p-Chk2 abundance in individual nuclei as in (**B**).

**E.** Comparison of relative yH2AX, pChk1, and pChk2 levels in cisplatin or etoposide-treated DTePs and CDK1i-induced endocycling cells, grouped by cell cycle phase. Data from 2 independent replicates (points) along with mean abundance (black bars).
